# The fitness cost of mismatch repair mutators in *Saccharomyces cerevisiae*: partitioning the mutational load

**DOI:** 10.1101/639765

**Authors:** Benjamin Galeota-Sprung, Breanna Guindon, Paul Sniegowski

## Abstract

Mutational load is the depression in a population’s mean fitness that results from the continual influx of deleterious mutations. Here, we directly estimate the mutational load in a population of haploid *Saccharomyces cerevisiae* that are deficient for mismatch repair. We partition the load in haploids into two components. To estimate the load due to nonlethal mutations, we measure the competitive fitness of hundreds of randomly selected clones from both mismatch repair-deficient and - proficient populations. Computation of the mean clone fitness for the mismatch repair-deficient strain permits an estimation of the nonlethal load, and the histogram of fitness provides an interesting visualization of a loaded population. In a separate experiment, in order to estimate the load due to lethal mutations (i.e. the lethal mutation rate), we manipulate thousands of individual pairs of mother and daughter cells and track their fates. These two approaches yield point estimates for the two contributors to load, and the addition of these estimates is nearly equal to the separately measured short-term competitive fitness deficit for the mismatch repair-deficient strain. This correspondence suggests that there is no need to invoke direct fitness effects to explain the fitness difference between mismatch repair-deficient and - proficient strains. Assays in diploids are consistent with deleterious mutations in diploids tending towards recessivity. These results enhance our understanding of mutational load, a central population genetics concept, and we discuss their implications for the evolution of mutation rates.

## Introduction

An evolving population experiences a continual influx of mutations, the vast majority of which, excluding neutral mutations, are likely to be deleterious (Fisher 1930). A deleterious allele in a haploid population will attain an equilibrium frequency that is the quotient of the mutation rate to that allele and the selection coefficient against it (Danforth 1923). The influx of deleterious mutations causes a depression in the population’s mean fitness that is termed the mutational load (Muller 1950), and the load at equilibrium is equal to the deleterious mutation rate (Haldane 1937). Because all populations experience mutation, all populations experience load, and a substantial proportion of the genetic variance for fitness in natural populations is due to mutational load (Charlesworth 2015).

Mutational load is closely connected to the evolution of mutation rates. Consider an asexual population in which there is genetic variation for the mutation rate: within such a population, distinct lineages with differing mutation rates experience differing loads and therefore possess differing mean fitnesses. In this way a downward selective pressure on the mutation rate is realized. This pressure is indirect in the sense that modifiers of the mutation rate are subject to selection without affecting any physiological property immediately related to fitness. The existence of ancient and highly conserved systems for replication fidelity (including proofreading, mismatch repair, and nucleotide excision repair) attests to the persistence of this selective pressure (Raynes and Sniegowski 2014).

In evolving populations, lineages with higher mutation rates (“mutators”) are continually produced by mutation to any of numerous mutation-rate-affecting loci. In the absence of beneficial mutations, the expected frequency of mutators within a population depends on the increase in the deleterious mutation rate caused by the mutator allele, the rate of mutation from wild type to mutator, and the mean selective effect of newly arising deleterious mutations (Johnson 1999; Desai and Fisher 2011). Investigations of natural and clinical isolates of *Escherichia coli* and other bacteria have shown that mutators of one to two orders of magnitude in strength, often defective in mismatch repair, are present at low but notable frequencies in many populations (Jyssum 1960; Gross and Siegel 1981; LeClerc et al. 1996; Matic et al. 1997; Oliver et al. 2000; Denamur et al. 2002; Richardson et al. 2002; Trong et al. 2005; Denamur and Matic 2006; Gould et al. 2007; reviewed in Raynes and Sniegowski 2014). Evolution experiments conducted with *E. coli* have demonstrated that mutators can displace wild types by virtue of their increased access to beneficial mutations (Cox and Gibson 1974; Chao and Cox 1983; Sniegowski et al. 1997; Giraud et al. 2001; Shaver et al. 2002; de Visser and Rozen 2006). Similar findings have been reported for *Saccharomyces cerevisiae* (Thompson et al. 2006; Raynes et al. 2011, 2018). However, in contrast to findings in prokaryotes, mismatch repair-deficient (henceforth *mmr*) or other types of strong mutators have not been found in natural *S. cerevisiae* populations (but see Bui et al. 2017; Raghavan et al. 2018), though weaker variation for the mutation rate has been detected (Gou et al. 2019). One explanation for this difference could be that *mmr* mutators experience higher load, compared to the wild-type, in *S. cerevisiae* than they do in *E. coli*. Indeed, it has been observed by several investigators that haploid *mmr S. cerevisiae* strains decline in frequency in the short term when co-cultured with wild-type strains (Thompson et al. 2006; Raynes et al. 2011, 2018; Bui et al. 2017), even if they eventually out-adapt the wild type. While this short-term deficit of the fitness of *mmr* mutators relative to the wild type has been attributed to increased mutational load in the *mmr* strain, the evidence that this is the case has been mostly circumstantial (but see Wloch et al. 2001) because it is generally difficult to rule out an additional direct fitness effect of any allele thought to cause an indirect fitness effect (Raynes and Sniegowski 2014).

In this work, we establish, by short-term competitive fitness assays and in agreement with prior studies, that an *mmr* haploid *S. cerevisiae* strain is substantially less fit than an otherwise isogenic *MMR+* (i.e. wild-type) strain. This fitness difference could be caused solely by load, or solely due to some direct fitness effect of the *mmr* phenotype; or it could be some combination of the two. We develop separate assays to measure the components of load due to nonlethal and lethal deleterious mutations. To estimate the load caused by nonlethal deleterious mutations, we randomly sampled hundreds of clones from *mmr* and wild-type populations and measured the competitive fitness of each. The resulting histogram of the distribution of fitness of the *mmr* population provides an illustration of the effect of a high mutation rate on population mean fitness. We find that the means of these distributions differ, indicating substantial load for the *mmr* strain, but not fully accounting for the total observed fitness difference between *mmr* and wild type strains. To estimate the lethal mutation rate under the two different mutational regimes, we manipulate single cells to track the fate of mother/daughter duos. We show that these two separately measured components of load— due to nonlethal and lethal mutations—approximately sum to the measured fitness difference between the strains; hence we find no reason to suppose a direct fitness effect for *mmr*. Investigations with diploid versions of our strains provide support for this conclusion. We discuss some implications of these findings for continued experimental and theoretical investigations into the evolution of mutation rates.

## Materials and Methods

### Data analysis and figure production

Data processing and analysis was performed in R (R Core Team 2019) and RStudio (RStudio Team 2015). Graphical output was produced using the package *ggplot2* (Wickham 2016).

### Strains

yJHK112, a haploid, prototrophic, heterothallic, MATa, BUD4-corrected, ymCherry-labeled W303 strain, was used as the haploid wild type in all work described in this paper. yJHK111, labelled with ymCitrine (a variant of YFP) and otherwise isogenic to yJHK112, was used as the “reference strain” in all haploid fitness competitions. These strains have been previously described (Koschwanez et al. 2013) and were generously provided by the laboratory of Andrew Murray, Harvard University, Cambridge, MA. An *msh*2*Δ* derivative of yJHK112, in which the wild-type *MSH2* allele was replaced with a kanMX geneticin (G418) resistance cassette (Wach et al. 1994), was used as the haploid *mmr* mutator strain in all work described in this paper. This strain was generously provided by Yevgeniy Raynes of the laboratory of Dan Weinreich, Brown University, Providence, RI and has been previously described (Raynes et al. 2018). The kanMX cassette has been shown to not have a negative effect on growth (Baganz et al. 1997; Goldstein and McCusker 1999).

We constructed diploid versions of each of the three above strains by transforming (Gietz and Schiestl 2007) each with plasmid pRY003, temporarily providing a functional HO locus allowing mating type switching and subsequent mating. pRY003 was a gift from John McCusker (Addgene plasmid #81043; *http://n2t.net/addgene:81043*; RRID:Addgene_81043). The diploid state of resulting isolates was confirmed by (1) ability to produce tetrads after plating to sporulation media; (2) by flow cytometry for total genomic content (following Gerstein et al. 2006); and (3) by the presence of a PCR product for both the *MATa* and *MATα* loci. The *mmr* diploids would not sporulate, but were confirmed to be diploids by the other two methods.

### Growth conditions

The liquid medium for all fitness competitions was synthetic dextrose (SD) minimal media containing yeast nitrogen base at a concentration of 6.7 g/L and glucose at a concentration of 1.5 g/L (0.15%), supplemented with tetracycline (15 mg/L) and ampicillin (100 mg/L). Fitness competitions were conducted in volumes of 200 ul media in deep polypropylene 96-well plates (Nunc 260251) sealed with flexible caps (Nunc 276002) and shaken at 1000 rpm with an orbit of 3mm (Corning LSE 6780-4) at a temperature of 30 C.

Initial growth in liquid for the lethal event assays was performed in SD as described above but without antibiotics, in flasks shaken at 200 rpm at 30C. Growth on agar SD (2% glucose, no antibiotics) plates for the lethal assays took place at room temperature, approximately 24 C.

### Competitive fitness assays and isolation of clones

Yeast, when grown by batch transfer with glucose as the carbon source, follow a relatively complex cycle of lag, fermentation, and respiration, and fitness benefits “accrued” in one phase (e.g. respiration) may not be “realized” until the next (e.g. the lag following the next transfer) (Li et al. 2018). Therefore we conducted short-term competitive fitness assays between wild-type and *mmr* genotypes in which strains were mixed for one growth cycle prior to measuring frequencies (essentially, following Gallet et al. (2012)). The fitness assays were conducted as follows, with the interval between each consecutive day spanning 24 hours. Day 1: wild-type, *mmr*, and the *YFP+* reference strain were inoculated from frozen stock into single wells. Day 2: each strain was transferred to a new well with fresh media, diluting 1/100. Day 3: competing strains were mixed 1:1 by volume and transferred to new wells with fresh media, diluting 1/100, to create between 6 to 8 replicate competitions. Day 4: competitions were transferred to new wells with fresh media, diluting 1/100, and the frequencies of the competitor and reference strain were assayed by flow cytometry (Guava EasyCyte). Discrimination between strains was performed on the SSC/GRN scatter plot. Day 5: the frequencies of the competitor and reference strain were again assayed by flow cytometry. The population density at the end of a 24-hour cycle was ∼2 × 10^7^ cells/mL; the census population size was thus ∼4 × 10^6^ at transfer and ∼4 × 10^4^ just after transfer.

The change in frequencies between Days 4 and 5 was used to calculate a selection coefficient *s*. Under a continuous model of growth (Crow and Kimura 1970 p. 193)

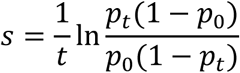

from which a relative fitness *w* = 1 + *s* follows. The number of generations, *t*, was assumed to be log_2_100, or approximately 6.64. *p*_0_ and *p*_*t*_ are the starting and ending frequencies of the genotype being tested (i.e. the frequencies at Days 4 and 5 in the above procedure). The resulting selection coefficients represent differences in Malthusian parameter (that is, the log of Wrightian fitness) scaled per generation of growth.

We conducted fitness competitions using this procedure in 8 separate blocks, each with multiple replicates as described above. Each block was begun on a different date. For each block, we computed the fitness of the *mmr* strain relative to the fitness of the wild-type strain by subtracting their mutual relative fitnesses to the reference strain. Each block included competitions in both haploid and diploid genotypes. Our final point estimate of the fitness difference between *mmr* and wild-type strains is the mean difference across all blocks, and the 95% confidence intervals (as shown in Fig. 2) were computed from the set of point estimates according to the t-distribution.

**Figure 1.**
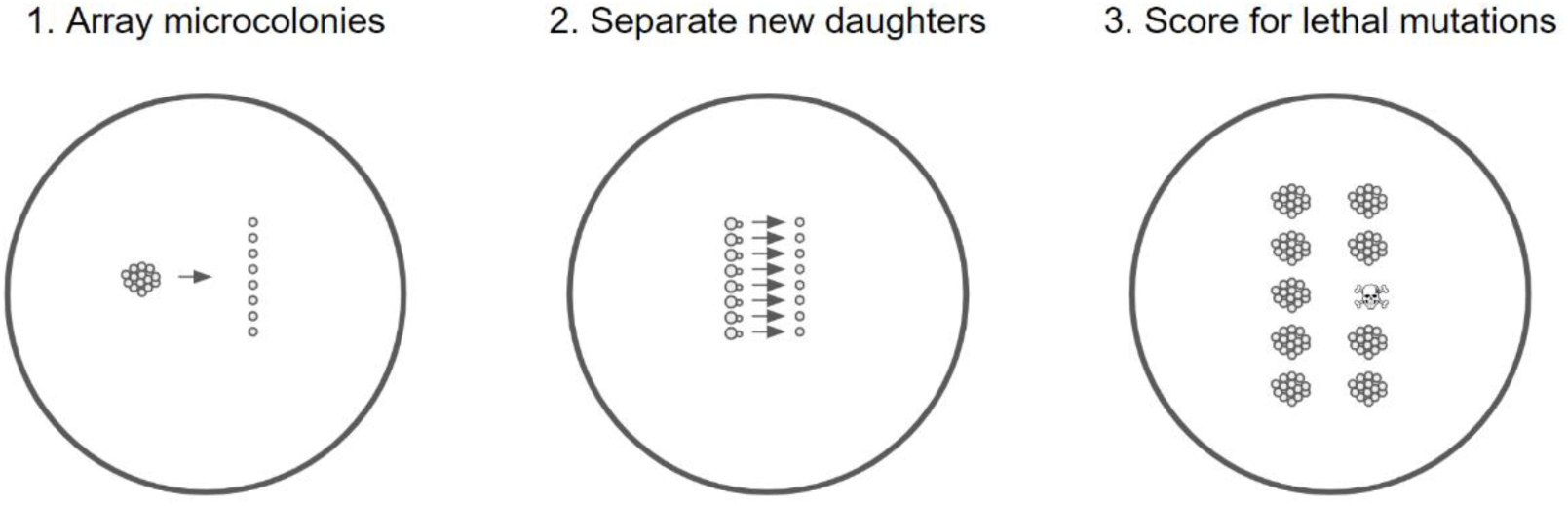
Schematic of lethal assay. The arraying in step 1 and separation in step 2 were performed by micromanipulation. Microcolonies in step 1 comprised on average 23 cells, and were founded by new daughter cells that had themselves been isolated by micromanipulation.

**Figure 2.**
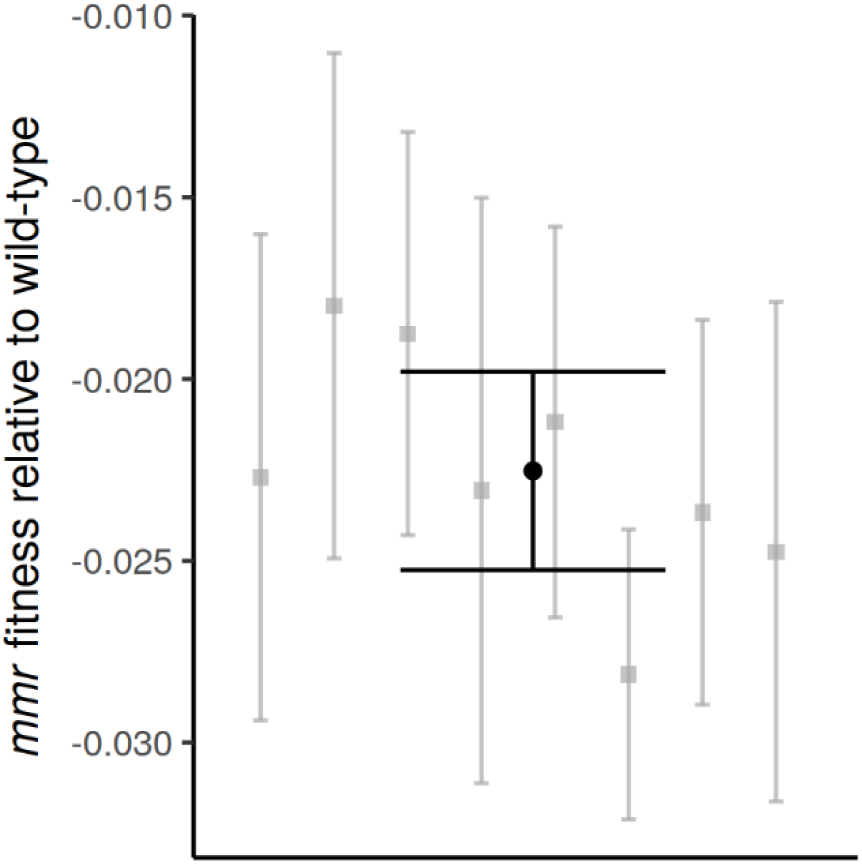
The average competitive fitness deficit (in black, error bars are 95% confidence intervals) of the mutator strain relative to the wild-type, expressed as selection coefficient, is 2.3%. Fitness competitions were conducted in a series of 8 blocks, shown in gray. The two strains were not competed directly against each other; within each block, each was competed separately against an otherwise isogenic *MMR+ YFP+* strain. Each block contained between 5 and 8 replicate competitions.

In 2 of the 8 fitness competition blocks, we randomly sampled individual clones. To do so, we additionally propagated the haploid wild-type and *mmr* strains on Day 3 in addition to mixing them 1:1 with the reference strain. Then, on Day 4, we plated these cultures, diluting appropriately, to YPD agar plates. After sufficient incubation, the resulting colonies were picked by pipet tip into wells containing 200 ul YPD, grown for 24 hours, and frozen down by mixing 1:1 with 30% glycerol and storing at −80 C until needed for fitness assays. The random selection of colonies was ensured by either (1) picking all colonies on a given plate or (2) picking colonies concentrically from a randomly placed point.

Fitness assays for sets of isolated clones were conducted on a single 96-well plate, which allowed us to assay the fitnesses of 88 clones (some wells being reserved for various purposes) or fewer per run. We followed essentially the same procedure as the 5-day competition described above, except that frequencies were estimated at Day 3 and Day 4 instead of Day 4 and Day 5. This modification was made because some clones had such reduced fitnesses that an extra day of growth after mixing 1:1 with the reference strain caused the starting frequency of the clone to depart too greatly from 50%.

The expected variance in measured fitness due to random sampling effects during flow cytometry was computed by means of a simple simulation in which the true frequency of each genotype at the start and end of the fitness competition was replaced by a random binomial variable. 10,000 replicates of this simulation were run.

### Calculation of nonlethal load

Mutational load is classically defined (Bürger 1998) as

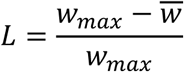

where 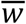 is the mean fitness of the population and *w*_*max*_ is the fitness of the fittest genotype.

We measured all fitnesses relative to a common fluorescent strain, as described above. We define the unloaded fitness of each genotype as equal to 1 and we expect no beneficial mutations to rise to appreciable frequency in the short course of these experiments. Hence, *w*_*max*_ = 1 and thus

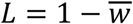

We expect measured selection coefficients to be distributed approximately normally around the true value, because of various sources of error including binomial sampling error, drift, instrument noise, environmental perturbations to individual wells (within-batch effects) and among 96-well plates (across-batch effects). To eliminate across-batch effects, for each run we adjusted all measured fitnesses by a constant *c* such that the mode fitness is 1 (equivalently, such that the mode selection coefficient is zero). Once this adjustment has been made,

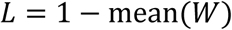

where *W* is the vector of all sampled clone fitnesses, or equivalently

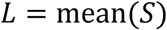

where *S* is the vector of all sampled clone selection coefficients. A 95% confidence interval for the nonlethal load was computed by bootstrapping from the measured fitnesses of all sampled clones: for 10,000 replicates, fitnesses were sampled randomly with replacement and the mean computed; from this empirical distribution, the 0.025 and 0.975 quantiles formed the bounds of the confidence interval.

### Lethal event assay

Strains were inoculated from frozen stock into 6 mL SD in a flask, and transferred to fresh media after 24 hours, diluting 1/100. After an overnight of growth, a streak from the culture was made onto an SD agar plate and five single cells with nascent buds were physically isolated by means of a Zeiss (West Germany) micromanipulating microscope fitted with a Singer Instruments (Somerset, UK) dissecting needle. These cells were periodically checked over the next few hours and the daughters physically separated once developed. These daughters became the founders of microcolonies that were allowed to grow at room temperature for ∼20 hours, reaching an average size of 23.3 cells (22.8 for the wild type, 23.8 for *mmr*, difference not significant at *p* > 0.6). These microcolonies were then dissected into a gridlike arrangement of single cells (step 1 in Fig. 1). These cells were then checked at intervals of one to two hours and daughters separated as soon as possible (step 2 in Fig. 1). The colonies resulting from these mother/daughter duos were checked at ∼24, ∼48, and in some cases ∼72 and ∼96 hours after separation. A lethal event was recorded when the growth of a mother or daughter lineage ceased. In such cases cessation of growth was sometimes immediate and sometimes occurred after a few generations. In the latter cases the growth was generally markedly slowed by the first observation. In a few cases, slow but unceasing growth was noted: these are presumed to be cases in which a strongly deleterious mutation occurred, though we stress that this assay was not designed to detect nonlethal deleterious mutations.

As described in the Discussion, the difference in the rate of lethal events between the wild type and *mmr* strains was used as the estimate for the lethal mutation rate for the *mmr* strain. A 95% confidence interval for this difference in rates was computed by the *prop.test* function in R (Newcombe 1998).

### Fluctuation assays

To measure the mutation rate to 5-fluoroorotic acid (5-FOA) resistance, we employed the following procedure. Strains of interest were inoculated into 10 mL YPD, grown in flasks with shaking at 30 C for ∼24 hours, and then transferred to 30 mL fresh YPD diluting such that ∼200 cells were passaged, in replicates of 5. After ∼48 hours of growth, each replicate was plated without dilution to SD + 5-FOA (1 g/L) agar plates to estimate density and absolute number of resistants, and plated with a 10^−5^ dilution to YPD agar plates to estimate total population density and absolute number. Plates were counted after ∼48 hours of growth and mutation rates were estimated using the maximum likelihood method of Gerrish (2008). For each round of fluctuation tests, we estimated mutation rates for both wild-type and *mmr* strains simultaneously in order to minimize the influence of any uncontrolled sources of variation.

### Homopolymers per gene

The per-base rate of homopolymeric runs of various lengths in *S. cerevisiae* coding regions were computed by a custom Python script. The *S. cerevisiae* S288C reference genome was downloaded from www.yeastgenome.org.

## Results

### Mutation rate elevation in *mmr* strain

To confirm that the *mmr* (*msh*2*Δ*) strain is a mutator, we conducted fluctuation tests using resistance to 5-FOA as the selectable phenotype. Averaged across replicate fluctuation tests, we found a 20.8-fold increase (95% CI: 13.4-to 28.3-fold) in the mutation rate for the *mmr* strain relative to the wild-type (Fig. S1). This is likely an underestimate of the effective genome-wide increase in the mutation rate because *mmr* mutators have a greatly elevated indel rate for homopolymeric runs (Lang et al. 2013), of which *URA3*, the main locus involved in this fluctuation test, is relatively devoid (Fig. S4).

### Fitness disadvantage of *mmr* compared to wild-type

We competed *mmr* and wild-type strains against a common YFP+ reference strain. Averaged across 8 separate blocks of fitness competitions, we found the mutator to be less fit than the wild-type, with an average fitness deficit, expressed as a selection coefficient per generation, of ∼2.25% (95% CI: 1.98% to 2.53%) (Fig. 2).

### Estimation of nonlethal load

We randomly sampled individual clones from both *mmr* and wild-type populations and measured the competitive fitness of each clone. The sampled fitness distributions are shown in Fig 3. The *mmr* strain’s fitness distribution has a prominent left tail of less fit individuals. We calculated the load as the difference between the mode and the mean fitness; this is approximately zero for the wild-type strain and ∼1.65% (95% CI: 2.19% to 1.08%) for the *mmr* strain. The difference in load between the two strains is significant (*p* < 10^−8^). The fitness distributions for the *mmr* and wild-type are significantly different in shape (Kolmogorov-Smirnov test; *p* < 10^−8^), while the fitness distribution for the wild-type strain is not significantly different from a normal distribution with the same mean and variance (Kolmogorov-Smirnov test; *p* > 0.05).

**Figure 3.**
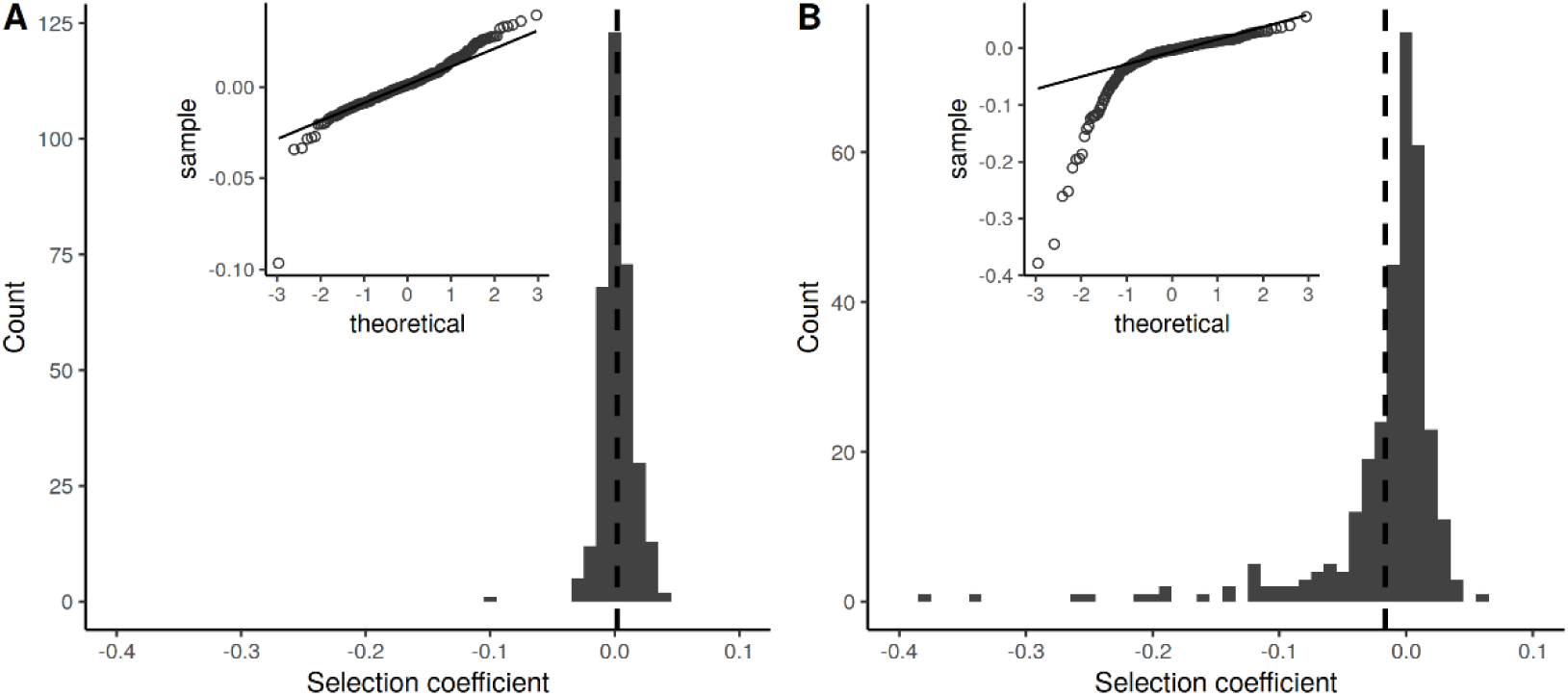
Distributions of fitness in haploid wild-types (**A**) and *mmr* mutators (**B**). We measured the fitnesses of 327 wild-type and 313 *mmr* clones. Fitnesses were measured in competition with an *MMR+ YFP+* reference strain otherwise isogenic to the wild-type, as described in Methods. Dashed vertical lines indicate the mean. The load is ∼0 for wild-types; for *mmr* mutators it is ∼1.7%. QQ plots of the fitness distributions are shown as insets. The two distributions differ significantly (Kolmogorov-Smirnov test, *p* < 10^−8^).

### Estimation of lethal mutation rate

To assay lethal mutation rates, we manipulated single *S. cerevisiae* cells, separating mother/daughter duos and tracking events in which one member of the duo failed to found a colony. The procedure is shown in Fig. 1 and explicated more fully in the methods. Assaying over 2200 duos for each strain, we found a rate of lethal events per newly replicated genome of 0.31% (95% CI: 0.018% to 0.055%) in the wild-type strain and 0.76% (95% CI: 0.53% to 1.07%) in the *mmr* strain (Table 1). The difference between these rates is 0.0044 (95% CI: 0.0012 to 0.0077). Because the observed rate of lethal events in the wild type is much higher than the expected lethal mutation rate, we take this difference as our estimate of the lethal mutation rate in the *mmr* strain (see the Discussion for elaboration on this point). Photographs of representative lethal events are shown in Fig. S3.

**Table 1.**

|  | Wild type |  | *mmr* |  |  |
| --- | --- | --- | --- | --- | --- |
| Event | Count | Rate | Count | Rate | p-value |
| Both OK | 2235 | 0.9967 | 2145 | 0.9908 | 0.0006 |
| One lethal | 14 | 0.0031 | 33 | 0.0076 | 0.0049 |
| One strongly reduced growth | 1 | 0.0002 | 7 | 0.0016 | 0.0363 |

Counts and frequencies of events of interest in the lethal assay. "Both OK" means that both mother and daughter cell grew into normal colonies. "One lethal" means that the lineage founded by either the mother or daughter cell ceased to grow within the observation period. "One strongly reduced growth" means that either the mother or daughter lineage was observed to grow noticeably slowly. Other events--both members of the duo lethal or strongly reduced growth, or the mother never budding--were not included in the analysis and are not displayed here. The p-values reflect the statistical significance of the difference in rates between wild type and *mmr* strains and were obtained by Fisher's exact test. Note that events are displayed per duo while rates are calculated per individual.

In our assay we followed separated duos that contained a suspected lethal until growth ceased. In some cases growth never ceased, but doubling times were very slow compared to the usual growth rate; such cases were not counted as lethal events but are tallied separately in Table 1. We also detected cases in which both members of a duo were lethal, or both showed strongly reduced growth, and also cases in which the mother cell never divided. Because such events probably stemmed from a mutation that occurred prior to the division that created the duo under observation, we excluded these events from our analysis.

## Results in diploids

From the haploid strains, we constructed diploid *mmr* and wild-type strains. We calculate the nonlethal load in the *mmr* diploid strain as ∼0 for the wild-type diploid strain and 0.30% (95% CI: −0.02% to 0.55%) (Fig S2) for the *mmr* strain—substantially less, by about 80%, than the equivalent load in the in *mmr* haploid strain (difference significant at *p* < 10^−4^). The diploid wild-type and *mmr* distributions differ significantly in shape (Kolmogorov-Smirnov test; *p* < 10^−4^) but do not differ significantly in mean (*p* > 0.05).

We also measured the difference in population fitness between wild-type and *mmr* diploid strains via short-term competitive fitness assays. We found that the *mmr* diploid is less fit than the wild-type diploid by a selection coefficient of ∼1.69% (95% CI: 1.40% to 1.94%) (Fig. 4). This difference, though larger than expected, is smaller than the fitness difference between wild-type and *mmr* haploid strains by 25% (haploid-to-diploid difference significant at *p* < 0.004).

**Figure 4.**
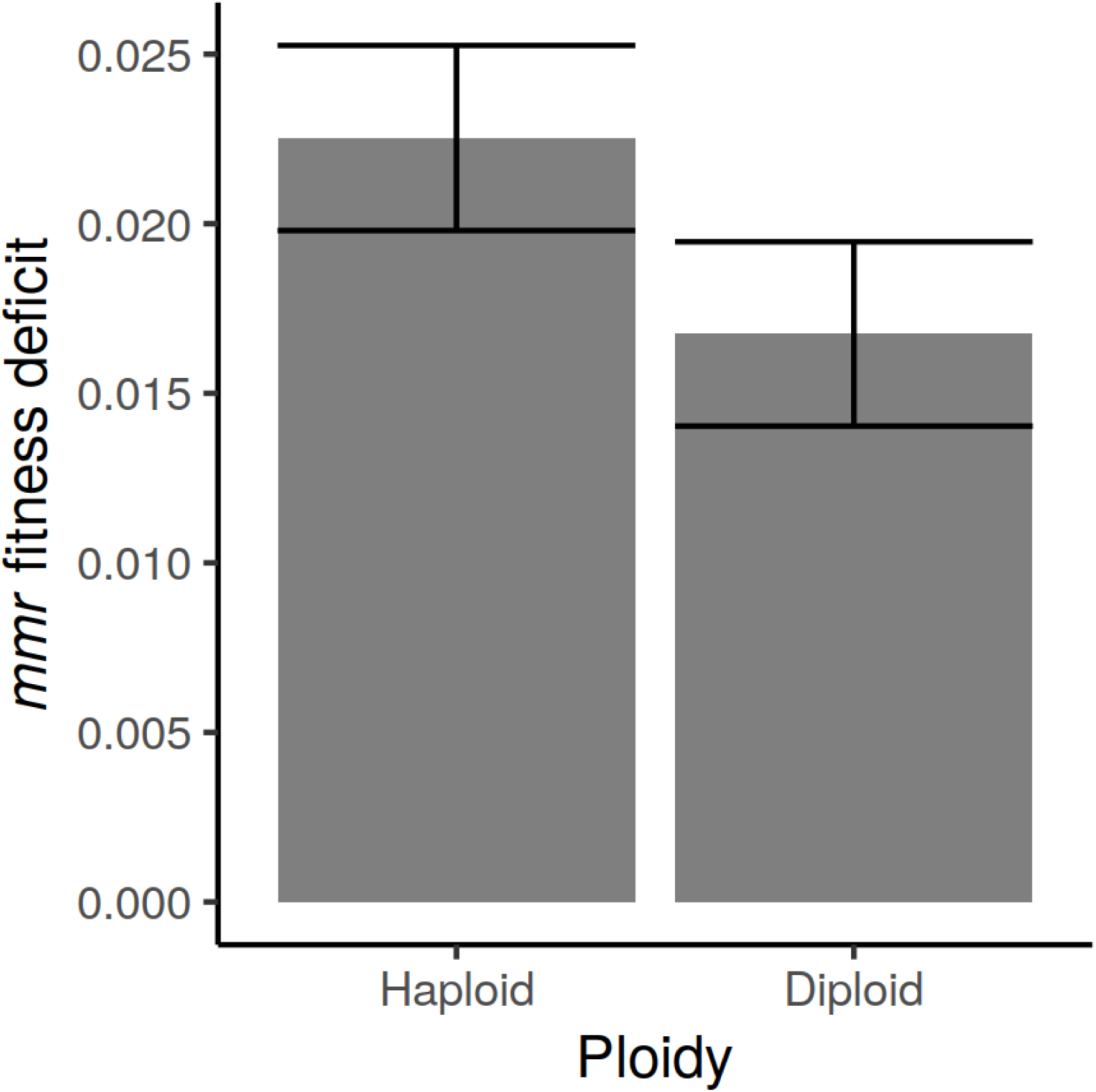
Change in *mmr* fitness disadvantage with ploidy. The fitness deficit, relative to the wild type, decreased by 26% from haploid to diploid strains (*p* < 0.004).

## Discussion

Prior work has shown that, over the short term, haploid *mmr S. cerevisiae* strains decline in frequency when competed with a strain that is wild-type for the mutation rate (Thompson et al. 2006; Raynes et al. 2011, 2018; Bui et al. 2017). Consistent with these findings, we find a fitness disadvantage, expressed as a selection coefficient per generation, of ∼2.3% for *mmr* haploids in short-term fitness competitions (Fig. 2). The magnitude of this selective disadvantage is similar to that in other reports, including Raynes et al. (2011) (2.4% cost), Raynes et al. (2018) (3.3% cost), and Wloch et al. (2001) (4.6% cost, though this is a noncompetitive measure of absolute growth rate).

The deleterious mutations that cause load include both lethal and non-lethal mutations. There is no fundamental theoretical distinction between these two classes of mutation insofar as their contribution to load is concerned: in many population genetic models, all members of an asexual population who are not of the least-loaded class are considered to be doomed (Rice 2002). However, their different manifestations require different experimental techniques. We therefore developed separate approaches to measure these two components of load.

### Load due to nonlethal mutations

We measured the short-term competitive fitnesses of 640 randomly selected haploid clones. The histogram and QQ plot for the haploid wild-type populations (Fig. 3A) suggest that, apart from one less-fit clone, the distribution of fitness for the wild-type strain is essentially normal. The normality of the distribution is consistent with nearly all wild-type clones having the same genotype and thus the same expected fitness, along with many small sources of error in estimation of fitness. One such source of error is drift over the course of the short-term fitness competition. The formula derived by Gallet et al. (2012) suggests that the expected variance in fitness measurement due to drift given our experimental parameters is ∼2 × 10^−6^. A larger source of variance is due to sampling error: in the fitness competitions, we estimate the relative frequencies of the competitors at two time points, sampling ∼8000 cells per time point. We carried out simulations that suggest that the expected variance due to sampling error is ∼2.3 × 10^−5^. These two sources of variance, summed, make up about 20% of the observed variance in selection coefficient. The remainder of the variance probably stems from small-scale environmental variation and other unknown sources of error.

In contrast to the results in wild types, the fitness histogram and QQ plot for the haploid *mmr* strain (Fig. 3B) are not reflective of a normal distribution. Instead, a prominent left tail of less fit clones demonstrates the effect of mutational load. The mean selection coefficient is ∼-1.7%, which is the quantification of the reduction in population mean fitness due to nonlethal load. This reduction accounts for a substantial portion (∼75%) of the measured competitive fitness difference (Fig. 2) between the two strains.

Our estimate of the nonlethal load (∼1.7%) reflects the average per-generation growth deficit of the mutator subpopulation due to the accumulation of deleterious mutations up to the point of random sampling of clones. We note that this is an estimate of the load at a nonequilibrium state, and is expected to be less than the full load achieved when mutation-selection balance is reached for all loci. Direct observation of mutation-selection equilibrium in a laboratory setting would be challenging because experimental populations rapidly generate adaptive mutations. Our strain-to-strain fitness assays, which found (Fig. 2) a fitness deficit of ∼2.3% for the *mmr* strain relative to the wild type, are likewise reflective of a nonequilibrium state. Since both estimates are derived from the same nonequilbrium populations, they are directly comparable.

Selection coefficients of about the magnitude we observe here cause changes in relative frequency that are extremely rapid in evolutionary terms. For example, a selective deficit of 2% would cause a decline from 50% frequency to 20% frequency in 70 generations. Observing a rapid initial decline of haploid *mmr S. cerevisiae* strains in competition with wild-types, some investigators (e.g. Grimberg and Zeyl 2005) have attributed the observed fitness difference to an unknown direct cost (i.e. a pleiotropic effect) while others (e.g. Raynes et al. 2018) have assumed that mutational load fully explains the dynamics. The question has remained open, in part because it has been nearly impossible to definitively rule out a direct fitness effect of being *mmr*—any attempt to measure such an effect will be confounded by the indirect fitness effects. By quantifying the indirect fitness effects (i.e. load) we seek to determine if a direct effect need be invoked to explain the observed experimental dynamics.

It is not surprising that the nonlethal load accounts for only a portion of the observed fitness difference. The nonlethal load assay relies on the growth of deleterious mutants in order to measure their fitness and thus cannot detect mutants who do not grow, i.e. lethal mutations. In order to measure this portion of the load, we designed an assay in which lethal events are directly observed.

### Load due to lethal mutations

The lethal mutation rate has long been a matter of interest (e.g. Dobzhansky and Wright 1941). By observing 4435 mother-daughter pairs (duos), we found a rate of lethal events of 0.0076 and 0.0031 for the *mmr* and wild-type strain, respectively.

Our observed wild-type lethal event rate, 0.0031, is on the order of estimates for the genomic mutation rate itself (Drake 1991; Lynch et al. 2008; Zhu et al. 2014; Sharp et al. 2018) and therefore cannot plausibly reflect the rate of lethal mutations. Our interpretation is that, for the wild-type, all or most observed lethal events were not caused by genomic mutations and are instead best considered to be non-mutational deaths, perhaps caused by fine-scale environmental fluctuations, experimental manipulation, or other stochastic sources of insult and stress. Observations of relatively high rates of cell death, too high to be due to lethal mutation, are not uncommon. Replicative aging studies of *S. cerevisiae* often observe low but substantial rates of cell death even in very young mother cells (e.g. Chiocchetti et al. 2007; Shcheprova et al. 2008). Rates of cell death on the order of our observed rate for the wild-type strain have also been observed in young bacterial cells (Wang et al. 2010), suggesting that relatively high rates of non-mutational, non-age-related deaths are common among microbes. Our assay design ensured that colonies were young (the oldest cell in a microcolony was on average ∼4.3 generations old) and we did not observe a bias in lethal events towards mothers (Table S1), so we do not attribute the observed lethal events to senescence. In fact, we observed, across both strains, a bias towards the lethal event occurring in the daughter cell. This difference was not statistically significant (*p* = 0.14), although within the *mmr* strain only we observed 10 lethal events in mothers and 23 in daughters (*p* = 0.04). The observed bias towards daughters dying, if not a sampling effect, could be attributable to smaller daughter cells being relatively more vulnerable to stress. Indeed, increased vulnerability of daughters to environmental sources of stress has been previously reported (Knorre et al. 2010).

An *a priori* estimation of the wild-type lethal mutation rate can be made as follows. Lang and Murray (2008) conducted careful estimations of the rate of loss-of-function mutations to the *CAN1* locus in a similar background (W303) as the strains used in this work. Multiplying this rate, 1.5 × 10^−7^, by the number of genes thought to be essential for viability, ∼1100 (Giaever et al. 2002), and accounting for the fact that *CAN1* is longer than the average essential gene gives an expected lethal rate in wild-type haploids of 1.5 × 10^−5^. This estimate is on the lower end but within the range of observed rates of accumulation of recessive lethals in several experiments conducted with diploids (Wloch et al. 2001; Hall and Joseph 2010; Nishant et al. 2010; Zhu et al. 2014; Jasmin and Lenormand 2016). Such a rate would suggest that we expected to observe about 0.3 lethal mutations in the wild-type strain in our experiment; we actually observed 14. Therefore, we consider the observed rate of lethal events in the wild-type to be an estimate of the rate of non-mutational deaths. The corresponding rate for the *mmr* strain is 0.0076 (difference significant at *p* < 0.001). Making the assumption that non-mutational deaths equally affect both strains, we take the difference between the wild-type and *mmr* lethal event rates, 0.0044 (95% CI: 0.0012 to 0.0077), as the estimate of the lethal mutation rate in the *mmr* strain. We note that our empirical result is fairly close to the figure obtained by multiplying the wild-type *a priori* estimate, 1.5 × 10^−5^, by the average fold increase in *CAN1* loss-of-function mutation rate for *mmr* strains in a collection of published reports (44-fold; see Table S4). A slightly different methodology, taking the average *CAN1* loss-of-function rate of *mmr* strains from published reports (1.5 × 10^−5^; Table S4) and multiplying by 1100 essential genes yields a somewhat higher expected lethal mutation rate of ∼0.015.

In many of the lethal events that we observed, growth did not immediately cease but continued for a few generations (Table S2) before halting. Limited growth for a few generations after an ultimately lethal mutation occurs has previously been observed (Mortimer 1955). We also observed morphological defects in several lethal events; one such instance is shown in the bottom panel of Fig. S3. We note that some lethal mutations that we observed could be due to chromosomal losses during mitosis (aneuploidies), but that knocking out *MSH2* has not been observed to greatly increase the rate of such events in haploids (Serero et al. 2014).

### Diploid findings

We measured the short-term competitive fitnesses of 573 randomly selected diploid clones. The distribution of fitness for *mmr* diploids (Fig. S2) suggests that they are substantially less loaded than *mmr* haploids, as would be expected if dominance attenuates the deleterious effects of new mutations. We calculate the nonlethal load in *mmr* diploids as ∼0.3%, as opposed to ∼1.7% in *mmr* haploids: that is, ∼80% of the load has gone away following diploidization. One interpretation of this finding is that deleterious mutations tend to be recessive in diploids. Thus, comparison of the fitness distributions of *mmr* diploids and *mmr* haploids is consistent with a high mutuation rate and diploidy shielding the effects of deleterious mutations.

The sampled wild-type diploid clones included more low-fitness individuals than the wild-type haploids (compare Fig. S2A to Fig. 3A). We cannot fully explain this observation; one possible explanation is that diploids are more prone than haploids to nondisjuctions causing aneuploid chromosomes, a notion for which there is some experimental support (Sharp et al. 2018).

The relative difference in short-term competitive fitness between wild-type and *mmr* strains is narrowed by 26% in diploids (Fig. 4). It is somewhat surprising, given that the nonlethal loads are not very different between wild-type and *mmr* diploids, that this figure is not larger. One possibility is that the diploid mutator fixed a deleterious mutation during the process of diploidization, which would account for the discrepancy between the reduction in nonlethal load (82%) and the reduction in total fitness difference (26%) in diploids compared to haploids. Another formal possibility is that diploid mutators have a higher lethal mutation rate than haploid mutators, but we cannot posit a causative mechanism for such an effect.

### Considering the two loads together

The total fitness difference between the haploid wild-type and *mmr* strains could be a consequence of greater mutational load for the *mmr* strain, a direct effect of the *msh*2*Δ* deletion, or a combination of the two. The addition of the lethal and nonlethal loads (0.0166 + 0.0044 = 0.0210) is approximately 7% smaller than the measured fitness difference (0.0225), and the difference is not significant (Fig. S5). The difference may simply be due to sampling error, or due to underestimation of one or the other of the loads. The nonlethal load may be slightly underestimated because clones were isolated by plating at the beginning of the growth cycle during which competitive fitness was measured. The load may have continued to increase somewhat during this growth cycle.

The broad equivalence of the sum of the loads, on the one hand, and the strain-to-strain competitive fitness, on the other, is consistent with the hypothesis that the total fitness difference is solely due to mutational load. Hence, although we cannot rule out the existence of a small direct fitness effect, these findings suggest that there is no need to invoke direct effects in explaining the fitness difference between the *mmr* and wild-type haploid strains.

The proportion of deleterious mutations that are lethal may also be inflated by an underestimation of the total deleterious mutation rate. The load is equal to the deleterious mutation rate only when the population is in mutation-selection balance. This equilibrium is reached instantly for lethal mutations, quickly for deleterious mutations of large effect, and very slowly for deleterious mutations of slight effect (Johnson 1999). The *mmr* populations in our assays experienced, including the initial process of transformation and growth before frozen storage, about 60 generations of growth, which is enough time to achieve mutation-selection balance for deleterious mutations of relatively large effect, but not for deleterious mutations of slight effect. Hence, our estimate of the total load (2.1-2.3%) should be considered an estimate of the lower limit for the deleterious mutation rate for *mmr* haploids.

### Comparison to results in bacteria

Insofar as *S. cerevisiae* and *E. coli* are two model organisms, from different domains of life, with which many evolution experiments have been performed, it is interesting to compare the loads of mismatch repair mutators in both. It appears that in *E. coli* the relative fitness deficit for *mmr* strains is smaller than it is in haploid *S. cerevisiae*. For instance, Shaver et al. (2002) did not detect a fitness difference between *mmr* and wild-type strains, de Visser and Rozen (2006) did not observe an initial decline in *mutS* frequency when that genotype was competed with the wild type at different starting ratios, and Boe et al. (2000) estimated at most a 1% selective disadvantage for *mmr* mutators. In this context it is relevant to note that there are several reports of *mmr* genotypes in natural *E. coli* isolates (LeClerc et al. 1996; Matic et al. 1997; Denamur et al. 2002), as well as in other types of bacteria (Oliver et al. 2000; Richardson et al. 2002; Trong et al. 2005; Gould et al. 2007). In *S. cerevisiae*, in contrast, no functionally *mmr* natural isolates have yet been found (though see Raghavan et al. 2018). Such observations suggest that *E. coli* may be relatively more robust than *S. cerevisiae* to the lack of a functional mismatch repair system. One reason for this difference could be that the genomic mutation rate in *E. coli* is lower than that of *S. cerevisiae* by a factor of about 4 (Lee et al. 2012, Lynch et al. (2008); Zhu et al. 2014; Sharp et al. 2018). If there is a similar absolute difference in deleterious mutation rate, then even if the relative fold increase in the deleterious mutation rate caused by the lack of mismatch repair is also equal in both organisms, the absolute difference in load, which is what controls the evolutionary dynamics, will be larger in *S. cerevisiae* than in *E. coli*. Another possible factor is differences in the spectrum and genomic substrate of mutations. In both *E. coli* and *S. cerevisiae*, the indel rate is greatly increased in *mmr* lineages, and the rate of indels is strongly elevated in homopolymeric repeats (HPRs). Both the relative increase from wild-type to *mmr* and the absolute indel rate in *mmr* are higher, and scale upwards faster with HPR length, in *S. cerevisiae* than in *E. coli* (Schaaper and Dunn 1991; Tran et al. 1997; Gragg et al. 2002; Lee et al. 2012; Lang et al. 2013). Examining all coding sequences in the *E. coli* and *S. cerevisiae* genomes, we find that there are significantly more homopolymeric repeats per coding genome, per gene, and per coding base in *S. cerevisiae* than in *E. coli* (Table S5). *S. cerevisiae* that are *mmr* are therefore relatively more burdened by indels than are *mmr E. coli* which could account for both the apparent larger fitness difference between *MMR+* and *mmr* and the corresponding apparent contrast in occurrence in natural isolates. We caution that this particular analysis is speculative in nature at this time: one important caveat is that, while this study and others have found large fitness differences between wild-type and *mmr* haploids, *S. cerevisiae* spend most of their time in nature as diploids, in which the fitness deficit of *mmr* lineages might be less severe: classically, equilbrium mutational load is halved in the recessive case compared to the additive, or haploid, case (Kimura et al. 1963). However, while estimates of the rate of outcrossing in *S. cerevisiae* are very low (Ruderfer et al. 2006), the rate of sporulation, which entails a haploid stage, is not known, and evidence of extensive inbreeding and loss of heterozygosity (Peter et al. 2018) suggest that it is relatively frequent. Recessive deleterious mutations may thus be frequently exposed to selection in natural *S. cerevisiae* populations by both the haploid life cycle stage and loss of heterozygosity from inbreeding, suggesting that diploidy may not be as much of a shield for *mmr* lineages as it otherwise would be. A second caveat is that, even if there is no direct fitness effect of *mmr* in haploids, there could be such an effect in diploids, perhaps due to misregulation of the frequency of recombination events (reviewed in Surtees et al. 2004; George and Alani 2012).

### Conclusions and future directions

We have found that the indirect fitness effects of strong modifiers for mutation rates are substantial in haploid *S. cerevisiae*, and that it is not necessary to postulate direct fitness effects in order to explain the selective disadvantage of the lack of a functional mismatch repair pathway. This finding is probably most relevant to experimental inquiries of the dynamics of mutation rate evolution in which *S. cerevisiae* is the model organism.

We have also reported findings relevant to fundamental questions about mutational dynamics, including the lethal mutation rate and the relative ratio of lethal and nonlethal deleterious mutations. By sampling the fitnesses of many individuals we have clearly demonstrated mutational load in an *mmr* population, and from the load we are able to estimate a lower limit for the deleterious mutation rate. We sampled hundreds of clones and were able to obtain a clear picture of the left tail of the fitness distribution for the *mmr* strain, but not for the wild-type strain. If the fitnesses of tens of thousands of clones could be measured, much could be learned about load and other evolutionary dynamics at wild-type mutation rates; such experiments may become possible as methods for high-throughput measurements continue to advance.

A limitation of this study is that we captured a snapshot of mutational load at a particular point in time in an evolving population. It would be interesting to observe, at a fine scale, how the distribution of fitnesses changes over time as a population approaches mutation-selection balance, adapts, and experiences other population genetic processes.

## Acknowledgments

This research was facilitated by a National Aeronautics and Space Administration grant (NNA15BB04A) to P.D.S. and a National Science Foundation Graduate Research Fellowship to B.G.S.

We thank the laboratories of Andrew Murray (Harvard University, Cambridge, MA) and Dan Weinreich (Brown University, Providence, RI) for their generous sharing of yeast strains.

## Data archiving

The authors plan to upload the raw data upon which this study is based to Dryad.

