## Supplemental figures and tables for "The fitness cost of mismatch repair mutators in *Saccharomyces cerevisiae*: partitioning the mutational load"

### Supplemental Figures

**Figure S1**
Four replicate fluctuation tests, using resistance to 5-fluoroorotic acid, show an average fold increase of 20.8 (95% CI: 13.4 to 28.3) for the *mmr* strain.

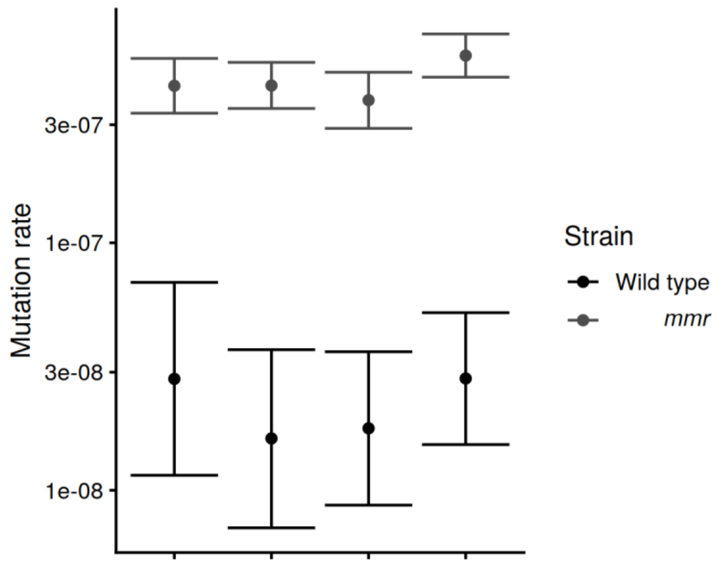

**Figure S2**
Histogram and QQ plot of fitness distributions for diploid strains. Panel **A** shows the results for the wild-type (n=279) strain and panel **B** for the *mmr* strain (n=294). As in Figure 3, the dashed line indicates the mean. The high fitness diploid *mmr* clone shaded in gray was removed from the data set for load calculations and is not shown on the histogram. With removal of this clone the load for the *mmr* diploids is 0.30%; without removal it is 0.28%.

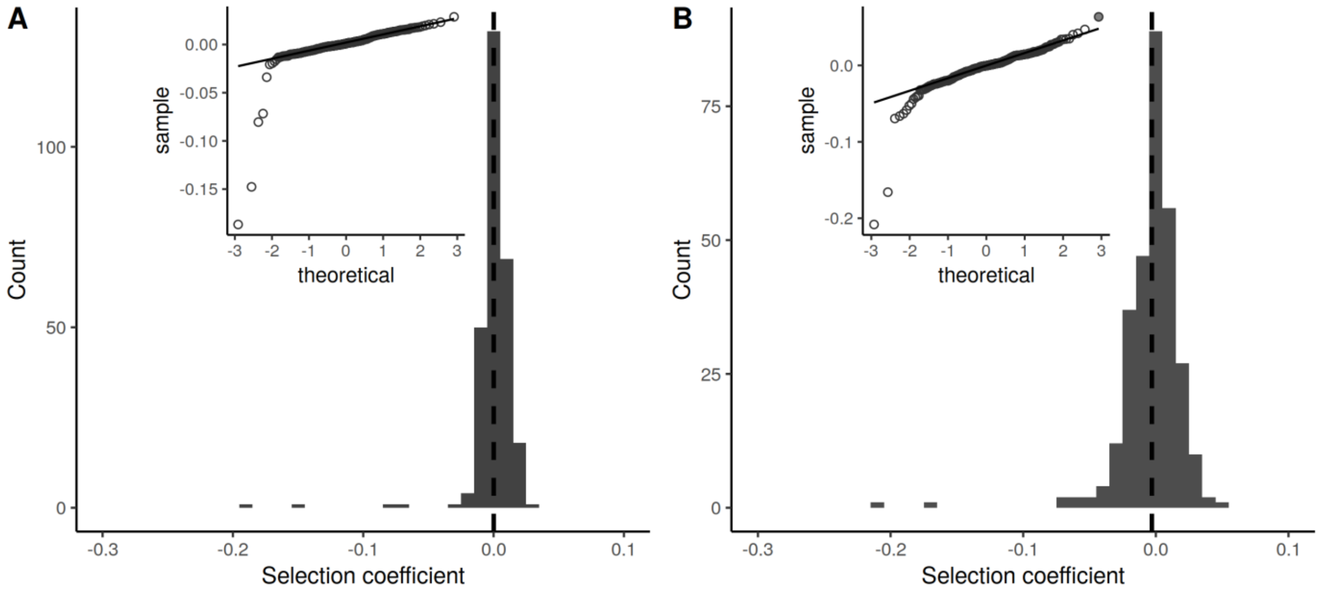

**Figure S3**
Photographs of two lethal events. Top: daughter is lethal, photo taken 27 hours after separation. Bottom: mother is lethal, photo taken 29 hours after separation. Scale bar is ~40 microns.

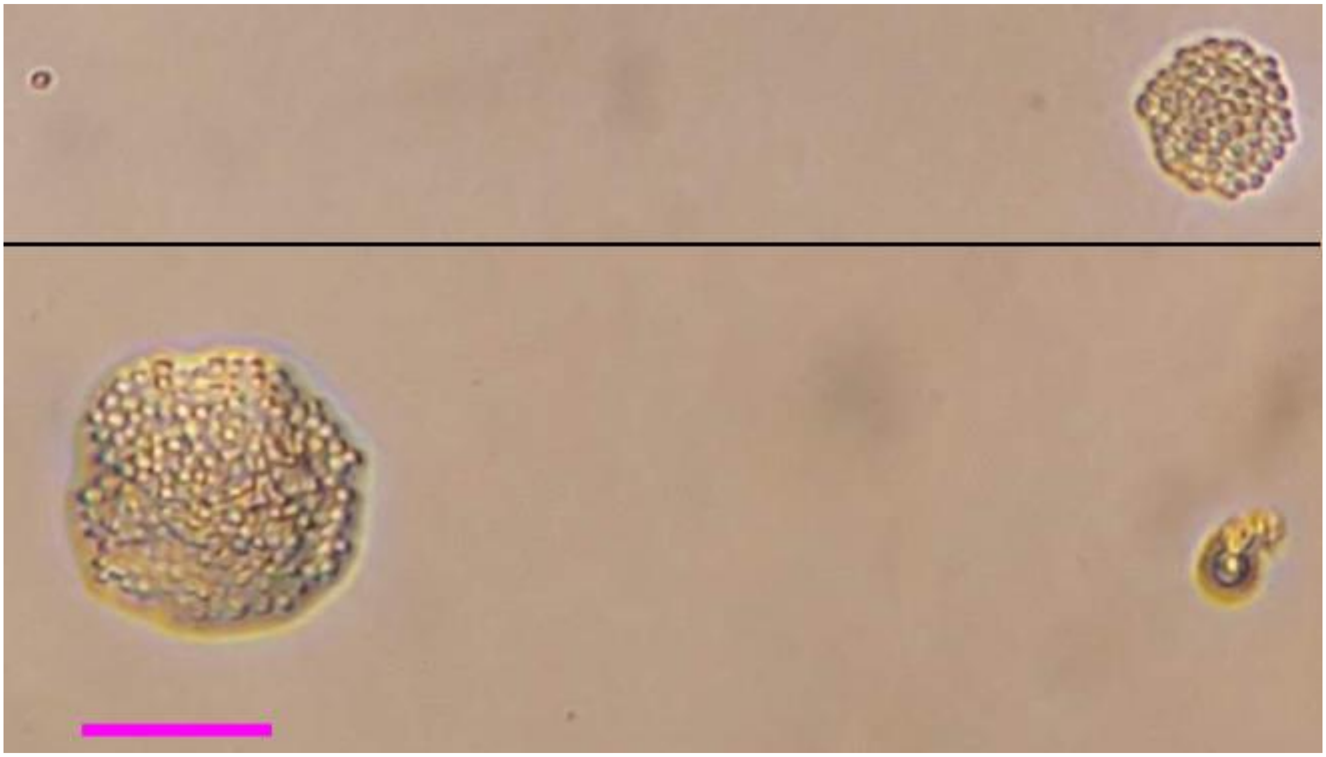

**Figure S4**
URA3 has fewer homopolymeric repeats than the “median gene”, the HPRs per gene as expected by the per-base HPR rate for the entire genome multiplied by the median gene length. This is largely but not entirely driven by URA3 being relatively short.

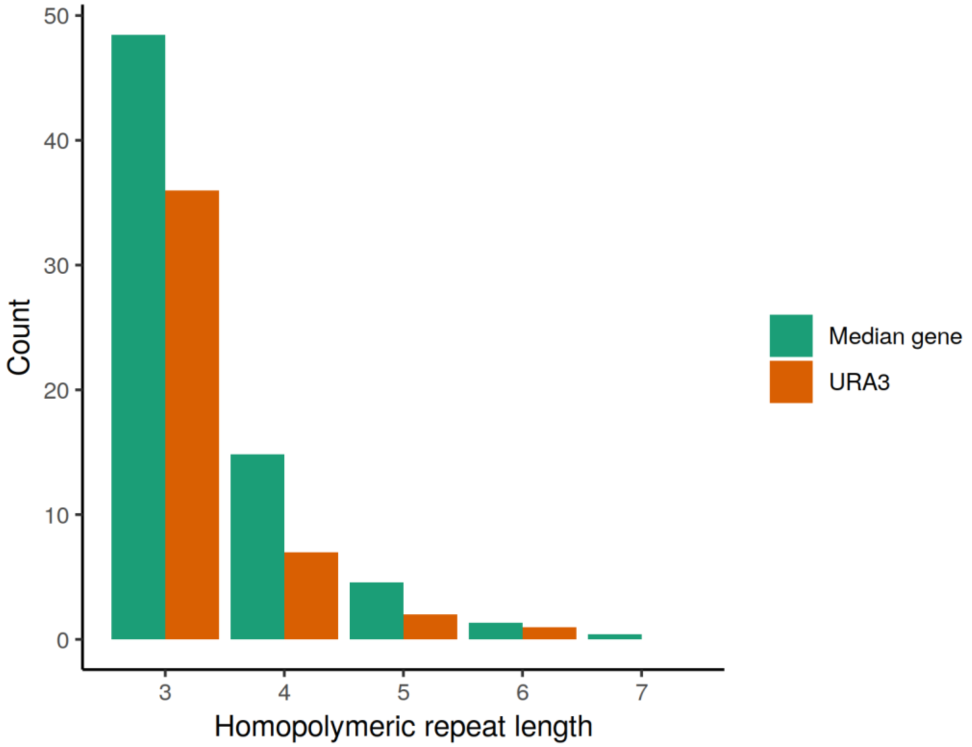

**Figure S5**
The sum of the nonlethal and lethal loads is ~93% of the measured competitive fitness deficit of the haploid *mmr* strain relative to the wild-type strain. The difference bewteen the two is not significant ($p > 0.2$). The difference between the competitive fitness difference and the nonlethal load alone is significant ($p < 0.02$).

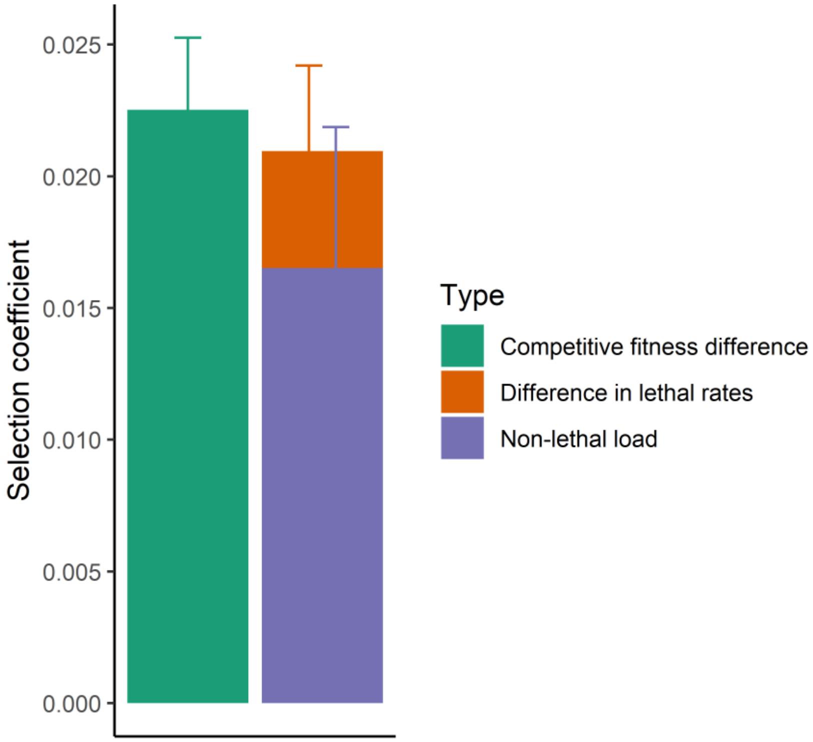

### Supplemental Tables

**Table S1:** Lethal events by mother/daughter.

| | Wild type | *mmr* | Total |
|---|---|---|---|
| Mother | 8 | 10 | 18 |
| Daughter | 6 | 23 | 29 |

**Table S2:** Lethal events by final microcolony size.

| Final size (number of cells) | Number |
|---|---|
| 1-2 | 15 |
| 3-30 | 25 |
| 31-100 | 3 |
| 100+ | 4 |
| Total | 47 |

**Table S3:** Lethal events by strain, size, and mother/daughter.

**Wild-type**

| Final size | Mother | Daughter | Total |
|---|---|---|---|
| 1-2 | 4 | 4 | 8 |
| 3-30 | 4 | 2 | 6 |
| 31-100 | 0 | 0 | 0 |
| 100+ | 0 | 0 | 0 |
| Total | 8 | 6 | 14 |

***mmr***

| Final size | Mother | Daughter | Total |
|---|---|---|---|
| 1-2 | 1 | 6 | 7 |
| 3-30 | 7 | 12 | 19 |
| 31-100 | 1 | 2 | 3 |
| 100+ | 1 | 3 | 4 |
| Total | 10 | 23 | 33 |

**Table S4:** Collected canavanine fluctuation tests for *S. cerevisiae.*

| Author and year | Strain | Locus | Genotype | Rate | Fold increase |
|---|---|---|---|---|---|
| Lang & Murray 2008 | W303 | CAN1 | WT | 1.5E-07 | n/a |
| Zeyl & de Visser 2001 | Y55 | CAN1 | WT | 3.2E-07 | |
| Zeyl & de Visser 2001 | Y55 | CAN1 | *msh2* | 1.7E-05 | 53 |
| Lang et al 2013 | W303 | CAN1 | WT | 8.0E-07 | |
| Lang et al 2013 | W303 | CAN1 | *msh2* | 6.7E-06 | 8 |
| Gammie et al 2007 | W303 | CAN1 | WT | 4.8E-07 | |
| Gammie et al 2007 | W303 | CAN1 | *msh2* | 1.5E-05 | 31 |
| Reenan & Kolodner 1992 | SK1 | CAN1 | WT | 4.0E-07 | |
| Reenan & Kolodner 1992 | SK1 | CAN1 | *msh2* | 3.4E-05 | 85 |
| Marsischky et al 1996 | MGD | CAN1 | WT | 1.0E-07 | |
| Marsischky et al 1996 | MGD | CAN1 | *msh2* | 4.0E-06 | 40 |
| Average of above | | CAN1 | WT | 3.8E-07 | |
| Average of above | | CAN1 | *msh2* | 1.5E-05 | 44 |

**Table S5:** Homopolymer occurrence in coding sequences of *S. cerevisiae* and *E. coli*. p-values are given for the difference in per base rates; "***" indicates a p-value of less than 2.2 x $10^{-16}$.

**Homopolymeric repeats in coding sequences**

| | *S. cerevisiae* | | | *E. coli* | | | *S. cerevisiae / E. coli* ratio | | | |
|---|---|---|---|---|---|---|---|---|---|---|
| Repeat length | Number | Per gene | Per base | Number | Per gene | Per base | Per coding genome | Per gene | Per base | p-value |
| 3 | 331178 | 57.31 | 4.0E-02 | 141596 | 32.830 | 3.5E-02 | 2.3 | 1.7 | 1.2 | *** |
| 4 | 102058 | 17.66 | 1.2E-02 | 35301 | 8.185 | 8.7E-03 | 2.9 | 2.2 | 1.4 | *** |
| 5 | 31363 | 5.43 | 3.8E-03 | 10895 | 2.526 | 2.7E-03 | 2.9 | 2.1 | 1.4 | *** |
| 6 | 9287 | 1.61 | 1.1E-03 | 2997 | 0.695 | 7.3E-04 | 3.1 | 2.3 | 1.5 | *** |
| 7 | 3066 | 0.53 | 3.7E-04 | 564 | 0.131 | 1.4E-04 | 5.4 | 4.1 | 2.7 | *** |
| 8 | 971 | 0.17 | 1.2E-04 | 97 | 0.022 | 2.4E-05 | 10.0 | 7.5 | 5.0 | *** |
| 9 | 306 | 0.05 | 3.7E-05 | 9 | 0.002 | 2.2E-06 | 34.0 | 25.4 | 17.0 | *** |
| ≥10 | 214 | 0.04 | 2.6E-05 | 0 | 0 | 0 | n/a | n/a | n/a | *** |
